# Epstein–Barr virus-encoded latent membrane protein 1 and B-cell growth transformation induces lipogenesis through fatty acid synthase

**DOI:** 10.1101/2019.12.15.876821

**Authors:** Michael Hulse, Sarah M Johnson, Sarah Boyle, Lisa Beatrice Caruso, Italo Tempera

## Abstract

Latent membrane protein 1 (LMP1) is the major transforming protein of Epstein-Barr virus (EBV) and is critical for EBV-induced B-cell transformation *in vitro*. Several B-cell malignancies are associated with latent LMP1-positive EBV infection, including Hodgkin’s and diffuse large B-cell lymphomas. We have previously reported that promotion of B cell proliferation by LMP1 coincided with an induction of aerobic glycolysis. To further examine LMP1-induced metabolic reprogramming in B cells, we ectopically expressed LMP1 in an EBV-negative Burkitt’s lymphoma (BL) cell line preceding a targeted metabolic analysis. This analysis revealed that the most significant LMP1-induced metabolic changes were to fatty acids. Significant changes to fatty acid levels were also found in primary B cells following EBV-mediated B-cell growth transformation.

Ectopic expression of LMP1 and EBV-mediated B-cell growth transformation induced fatty acid synthase (FASN) and increased lipid droplet formation. FASN is a crucial lipogenic enzyme responsible for *de novo* biogenesis of fatty acids in transformed cells. Furthermore, inhibition of lipogenesis caused preferential killing of LMP1-expressing B cells and significantly hindered EBV immortalization of primary B-cells. Finally, our investigation also found that USP2a, a ubiquitin-specific protease, is significantly increased in LMP1-positive BL cells and mediates FASN stability. Our findings demonstrate that ectopic expression of LMP1 and EBV-mediated B-cell growth transformation leads to induction of FASN, fatty acids and lipid droplet formation, possibly pointing to a reliance on lipogenesis. Therefore, the use of lipogenesis inhibitors could potentially be used in the treatment of LMP1+ EBV associated malignancies by targeting a LMP1-specific dependency on lipogenesis.

**Importance:** Despite many attempts to develop novel therapies, EBV-specific therapies currently remain largely investigational and EBV-associated malignancies are often associated with a worse prognosis. Therefore, there is a clear demand for EBV-specific therapies for both prevention and treatment of viral-associated malignancies. Non-cancerous cells preferentially obtain fatty acids from dietary sources whereas cancer cells will often produce fatty acids themselves by *de novo* lipogenesis, often becoming dependent on the pathway for cell survival and proliferation. LMP1 and EBV-mediated B-cell growth transformation leads to induction of FASN, a key enzyme responsible for the catalysis of endogenous fatty acids. Preferential killing of LMP1-expressing B cells following inhibition of FASN suggests that targeting LMP-induced lipogenesis could be an effective strategy in treating LMP1-positive EBV-associated malignancies. Importantly, targeting unique metabolic perturbations induced by EBV could be a way to explicitly target EBV-positive malignancies and distinguish their treatment from EBV-negative counterparts.

## Introduction

The Epstein-Barr virus (EBV) is a double-stranded DNA human gammaherpesvirus that latently infects approximately 95% of the population worldwide (1). EBV was the first human tumor virus identified (2) and contributes to about 1.5% of all cases of human cancer worldwide (3). Latent membrane protein 1 (LMP1) is expressed in the majority of EBV-positive cancers, including: Hodgkin’s and diffuse large B-cell lymphomas, HIV and post-transplant lymphoproliferative disorders, as well as nasopharyngeal and gastric carcinomas (11). *In vitro*, EBV is able to convert primary B-cells into immortalized lymphoblastoid cell lines (LCLs), and the EBV oncoprotein LMP1 is critical for this process (4, 5). LMP1 is a transmembrane protein containing two signaling domains: C-terminal-activating region 1 and 2 (CTAR1 and CTAR2). Through these two domains, LMP1 can mimic CD40 signaling to activate nuclear factor-κB (NF-κB), phosphoinositide 3-kinase (PI3K)/AKT, and Ras – extracellular signal-regulated kinase (ERK) – mitogen-activated protein kinase (MAPK) pathways (6). The activation of these signaling pathways by LMP1 contribute to its ability to transform cells by altering the expression of a wide range of host gene targets (7). LMP1 has also been shown to promote aerobic glycolysis and metabolic reprogramming in B cell lymphomas and nasopharyngeal epithelial cells (8–14). The transition from a resting B-cell to a rapidly proliferating cell following EBV infection, and the presence of EBV in associated malignancies, entails major metabolic changes. The role of LMP1 in these processes is incompletely understood. To further examine LMP1-induced metabolic reprogramming in B cells, we ectopically expressed LMP1 in an EBV-negative Burkitt’s lymphoma cell line preceding a targeted relative quantitation of approximately 200 polar metabolites spanning 32 different classes. The top metabolites induced by LMP1 were fatty acids. In parallel, the same metabolic analysis was carried out to compare metabolic changes in primary B cells following EBV-mediated B-cell growth transformation, which also revealed large changes in fatty acid levels.

Aerobic glycolysis is a well-established phenotype in cancer cells, and even though deregulated lipid metabolism has received less attention, it is just as ubiquitous as a hallmark of cancer (15). Non-transformed cells will preferentially obtain fatty acids from dietary sources for their metabolic needs versus *de novo* lipid synthesis (lipogenesis). However, despite access to these same dietary sources, cancer cells will often preferentially rely on endogenous fatty acids produced by *de novo* lipogenesis, often becoming dependent on the pathway for cell survival and proliferation. Fatty acids are essential for these processes as they are used as substrates for oxidation and energy production, membrane synthesis, energy storage and production of signaling molecules. Fatty acid synthase (FASN) is responsible for the catalysis of endogenous fatty acids and therefore is commonly upregulated in cancer cells (16–18). FASN condenses malonyl-CoA with acetyl-CoA, using NADPH as a reducing equivalent, to generate the 16-carbon fatty acid palmitate (19). In addition, upregulated glycolysis has been suggested as a mechanism for generating intermediates for fatty acid synthesis (20, 21). Once fatty acids are made, they can be converted to triglycerides and stored as lipid droplets for cellular energy storage (16). Lipid droplets can also contain phospholipids and sterols for membrane production (22).

There are two main pathways that transformed cells use to upregulate FASN, found at the levels of both transcription and post-translation. In the first case, FASN expression can be stimulated by the transcription factor sterol regulatory element-binding protein 1c (SREBP1c), which binds to and activates sterol regulatory elements (SREs) in the promoter region of FASN and other genes involved in lipogenesis (23, 24). SREBP1c is an isoform of the SREBF1 gene, which transcribes the two splice variants, SREBP-1a and SREBP-1c, that are encoded from alternative promoters and differ in their NH2-terminal domains (25). At the post-translational level, increased FASN protein levels can be obtained through interaction with ubiquitin-specific peptidase 2a (USP2a), a ubiquitin-specific protease that can stabilize FASN by removing ubiquitin from the enzyme (26). These two main methods of FASN regulation do not have to be mutually exclusive, it is also possible that they concurrently take place in cancer cells.

In this study, we determined that ectopic expression of LMP1 and EBV-mediated B-cell growth transformation leads to induction of FASN, fatty acids and lipid droplet formation. This points to a potential reliance on lipogenesis as demonstrated by preferential killing of LMP1-expressing B cells following inhibition of lipogenesis. It is therefore conceivable that use of lipogenesis inhibitors could play a role in the treatment of LMP1+ EBV associated malignancies by targeting LMP-induced metabolic dependencies.

## Results

### Fatty acids are the top metabolites increased by LMP1

To identify cellular metabolites that can be altered by LMP1, we first ectopically expressed LMP1 in the EBV-negative Burkitt’s lymphoma (BL) cell line DG75. Cells were transduced with retro-viral particles containing either pBABE-HA (empty vector) or pBABE-HA-LMP1 (LMP1) vectors as described previously (9). Using this cell system, we then undertook a targeted approach to determine the relative quantities of approximately 200 polar metabolites spanning 32 different classes to examine LMP1-induced metabolic changes. These changes are summarized by heat map and principal component analysis (PCA) (**Fig. 1A and 1B**). The unsupervised hierarchical clustering classified each sample groups into distinct clusters, indicating that LMP1+ cells possess a distinct metabolic profile compared to LMP1-cells (**Fig. 1A**). We observed a similar separation for the sample groups in the PCA analysis (**Fig. 1B**). However, the PCA analysis showed that the LMP1-samples do not completely cluster together. The lack of complete clustering in the PCA analysis is probably due to a few metabolites with much higher levels in the one of the LMP1-samples (pBABE untreated sample 1) skewing the PCA analysis. Nevertheless, our metabolic analysis indicate that distinct metabolic profiles exist between LMP1+ B-cells and LMP1-cells. To characterize the specific metabolites that are affected by LMP1, we further explored the data generated by our analysis that used mass spectrometry followed by hydrophilic interaction chromatography (HILAC). Peak areas, representing metabolite levels, were extracted using ThermoScientific Compound Discoverer 3.0. Metabolites were identified from a provided mass list, and by MS/MS fragmentation of each metabolite followed by searching the mzCloud database. Significant differences (q-value < 0.05) in proteins of least 1.5-fold between empty vector (pBABE) and LMP1 conditions (based on average value of the triplicate sample) were indicated as ‘True’ changes (**supplementary table 1**). When comparing pBABE vs LMP1 cell lines and sorting fold change of metabolites in descending order, the top 13 ‘True’ metabolites (confirmed using pure compounds) induced by LMP1 were fatty acids. These fatty acids were largely saturated medium and long chain and were increased from 2.64 to 36.42-fold change (**Fig. 1C**). Previously, we have shown Poly(ADP-Ribose) Polymerase 1 to be important in LMP1-induced aerobic glycolysis and accelerated cellular proliferation using the PARP inhibitor olaparib (9). Therefore, we included an olaparib treatment group in our metabolic analysis to examine whether PARP inhibition could offset LMP1-induced changes to cellular metabolites. Unsupervised clustering analysis and PCA analysis showed that metabolic changes induced by LMP1 expression could be partially reverted by treatment with the PARP inhibitor Olaparib (**Fig. 1A**). When we sorted the fold change of metabolites between pBABE vs LMP1 in descending order as described above, we found a nearly perfect inverse correlation between the fatty acids in our LMP1 untreated vs LMP1 + olaparib groups. In other words, 11 of the 13 fatty acids fatty acids that were most increased with ectopic expression of LMP1 were also the most decreased when these same cells were then treated with olaparib. Significant fold changes were in the range of −1.89 to −3.64 (**Fig. 1C**). This may partly explain the ability of olaparib to blunt the proliferative advantage bestowed by LMP1 that we previously reported (9). Finally, in a comparison of LMP1+ DG75s treated with olaparib compared to untreated pBABE, we found that each metabolite’s fold changes are roughly 50% less than those observed in LMP1+ untreated cells compared to pBABE. These results indicate that PARP inhibition offsets LMP1+ effects on cell metabolism.

**Figure 1.**
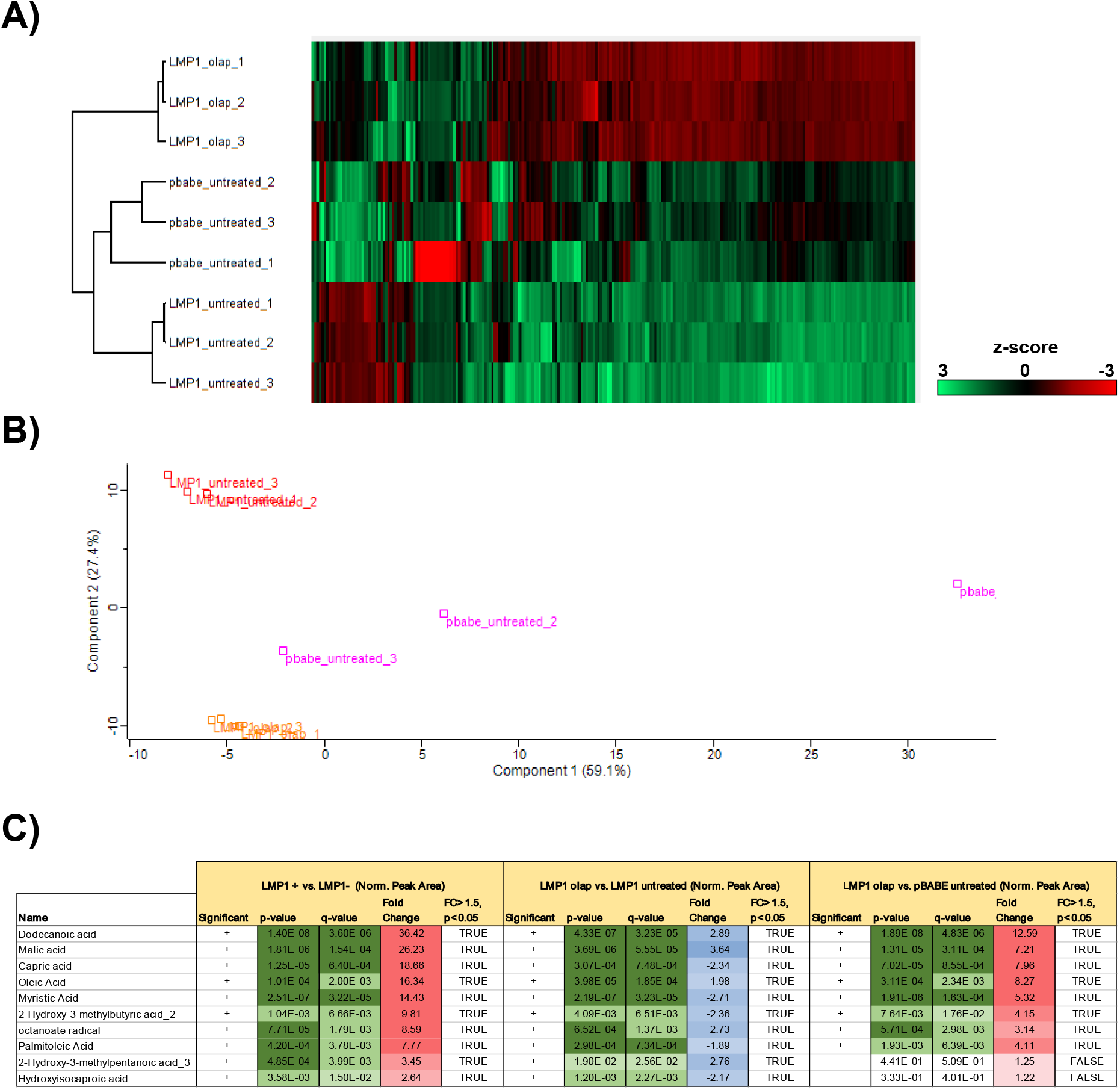
A targeted relative quantitation of approximately 200 polar metabolites spanning 32 different classes revealed fatty acids as the top metabolites induced by LMP1. **A)** Heat map comparing metabolite levels in DG75 transduced with retroviral particles containing either pBABE (empty vector) or pBABE-HA-LMP1 vectors. LMP1 + cells were incubated for 72 hrs with 2.5 μM olaparib or the DMSO vehicle as a control. Heat maps were generated using Perseus software by performing hierarchical clustering on Z-score normalized values using default settings (row and column trees, Euclidean distances, k-means preprocessing with 300 clusters). **B)** Principal component analysis (PCA), performed using default settings on Perseus software, of untreated LMP1+ and LMP1-cells and LMP1+ cells treated with olaparib. **C**) Peak areas, representing metabolite levels, were extracted using ThermoScientific Compound Discoverer 3.0. The peak areas were normalized using constant sum. Metabolites were identified from a provided mass list, and by MS/MS fragmentation of each metabolite follow by searching the mzCloud database (www.mzcloud.org). Comparisons between the conditions were performed: Student’s T-test p-value; q-value: Benjamini-Hochberg FDR adjusted p-value to account for multiple testing. q-value < 0.05 is considered significant and flagged with “+” in the “Significant” column; Fold change between 2 conditions (based on average value of the quadruplicate sample); Proteins displaying significant change (q-value < 0.05) with at least 1.5 fold change are indicated in the “FC>1.5, p<p0.05” column.

### LMP1 induces FASN and lipogenesis

Because fatty acids were the dominate metabolite class induced by LMP1, we sought to pursue a potential enzyme responsible. FASN catalyzes *de novo* lipogenesis and is commonly upregulated across many different cancers (16–18). Furthermore, a recent study demonstrated that LMP1 upregulates FASN and lipogenesis in EBV-positive nasopharyngeal carcinoma (NPC) (27). We therefore wanted to determine if LMP1 could induce FASN and lipogenesis in B-cells. Using western blotting, we showed that ectopic expression of LMP1 increased FASN protein levels around 2.5-fold as compared to empty vector control (**Fig. 2A and 2B**). To determine if the LMP1-mediated increase of fatty acids and FASN levels were inducing lipogenesis, we employed Nile Red staining, a potent and specific lipid droplet stain. Lipid droplets are small cytoplasmic organelles that can store fatty acids, providing available energy as well as cellular membrane material (16). Under serum-deprived conditions, we stained pBABE and LMP1 cells with Nile red followed by FACS analysis. We found that LMP1 led to an increase in Nile Red staining (**Fig. 2C**), which was then further quantified using a fluorescent plate reader (**Fig. 2D**). The somewhat modest increases in FASN and lipid droplet formation should be viewed in the context of the BL background used for the ectopic expression of LMP1, as alteration in lipid metabolism is a notable feature of BL and likely blunted the effect of LMP1 ectopic expression (28).

**Figure 2.**
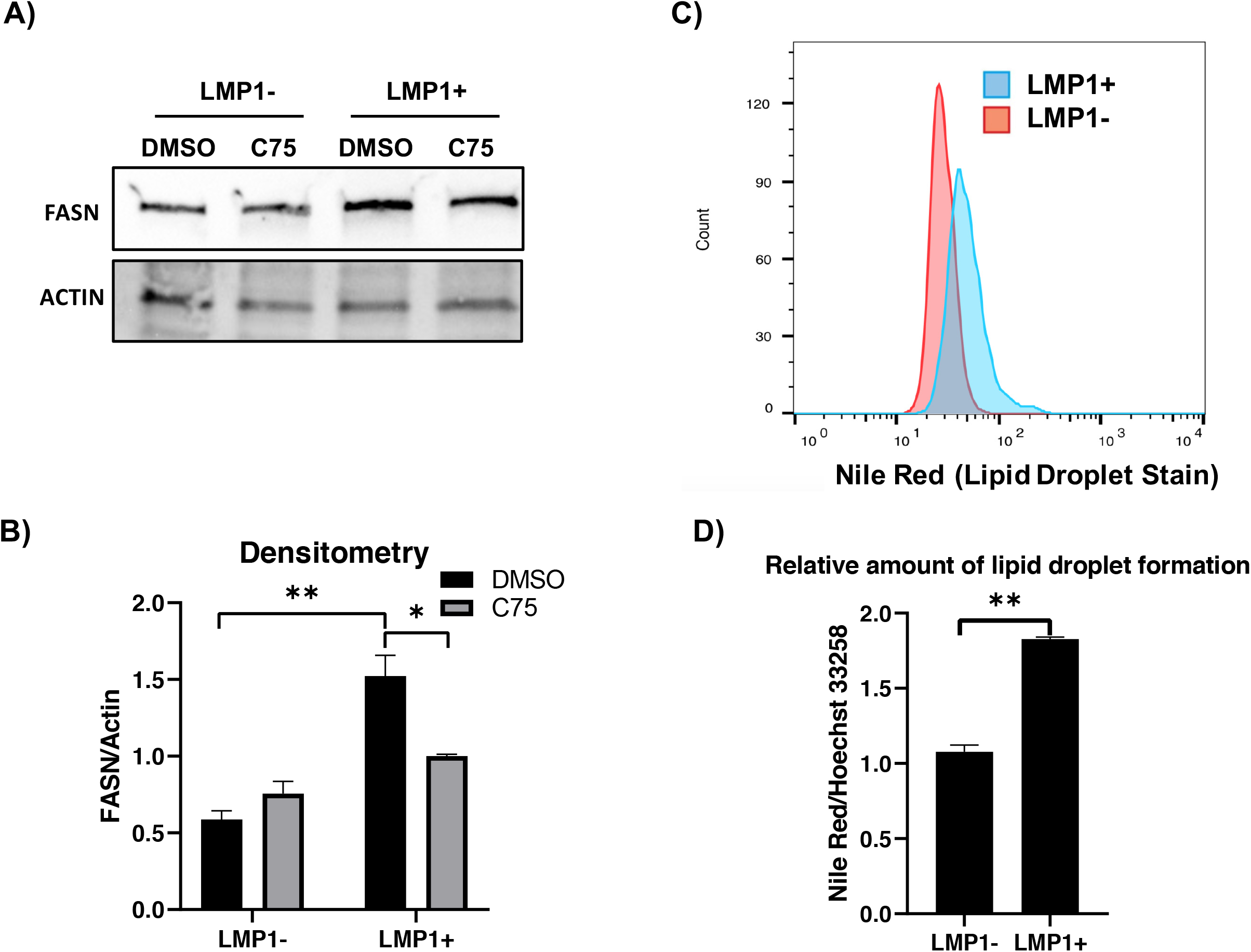
LMP1 leads to increased FASN and lipid droplet formation. **A)** Western blot of the EBV-negative B cell line DG75 transduced with retroviral particles containing either pBABE (empty vector) or pBABE-HA-LMP1 vectors and treated with 10 μg/mL of the FASN inhibitor C75 for 24 hrs. Cell lines were probed for FASN. Actin served as a loading control. **B)** Densitometry of FASN/Actin normalized to untreated empty vector (pBABE). **C)** FACs analysis of Nile Red fluorescence staining (excitation, 385 nm; emission,535 nm) for lipid droplets in DG75 cell line transfected with an empty plasmid vector or LMP1 expression construct. **D)** The relative amount of lipid droplet formation was calculated by plate reader by normalizing the Hoechst 33342 fluorescence (excitation, 355 nm; emission, 460 nm) to the Nile Red signal in each well. Error bars represent standard deviation of two independent experiments. P values for significant differences (Student’s t-test) are summarized by two asterisks (p<0.01) or one asterisk (p<0.05).

### EBV-immortalization of B cells leads to significant increases in metabolic cofactors and fatty acids

Our initial analysis into LMP1-mediated metabolic changes revealed that fatty acids were the major metabolites increased. However, we wanted to extend our examination of LMP1’s role in metabolic remodeling of the cell in the broader context of EBV-immortalization of B cells. To do this, using the same metabolic analysis as described for ectopic expression of LMP1, we infected primary B cells with EBV, resulting in their transformation into LCLs, a process in which LMP1 is critical (4, 5). Both primary B cells and their corresponding matched LCLs (60 dpi) were extracted for metabolite analysis. These changes are summarized by heat map and principal component analysis (PCA) demonstrating that EBV infected cells have a different metabolic profile compared to uninfected primary B cells (**Fig. 3A and 3B**). Interestingly, the highest metabolites induced (50-70-fold change) following immortalization of B cells was nicotinamide (NAM), nicotinic acid and nicotinamide adenine dinucleotide (NAD) (**Fig. 3C**). NAM and nicotinic acid are both precursors of NAD and nicotinamide adenine dinucleotide phosphate (NADP), which are both coenzymes in wide-ranging enzymatic oxidation-reduction reactions, including glycolysis, the citric acid cycle, and the electron transport chain (29). Of note, the reduced form of NADP, NADPH, is the critical reducing equivalent used by FASN to synthesize long chain fatty acids (30). NAD+ is also an essential cofactor for Poly(ADP-Ribose) Polymerase 1 (31) which we have previously shown to be important in EBV latency status and LMP1-mediated host gene activation (9, 32). Aside from these important metabolic cofactors, our metabolic analysis also revealed several fatty acids amongst the top metabolites induced following EBV transformation. These increases were in the range of 3-20-fold change and were mainly in the class of long and very long chain polyunsaturated fatty acids, differing form our ectopic LMP1 analysis where the top fatty acids were mainly saturated and medium to long chain length (**Fig 3C**). The differences observed in the fatty acid species between figure 3C and figure 1C are unsurprising as the DG75 established Burkitt’s lymphoma cell line most likely shifts the metabolic profile that would be observed in primary B-cells and their matched LCLs (28).

**Figure 3.**
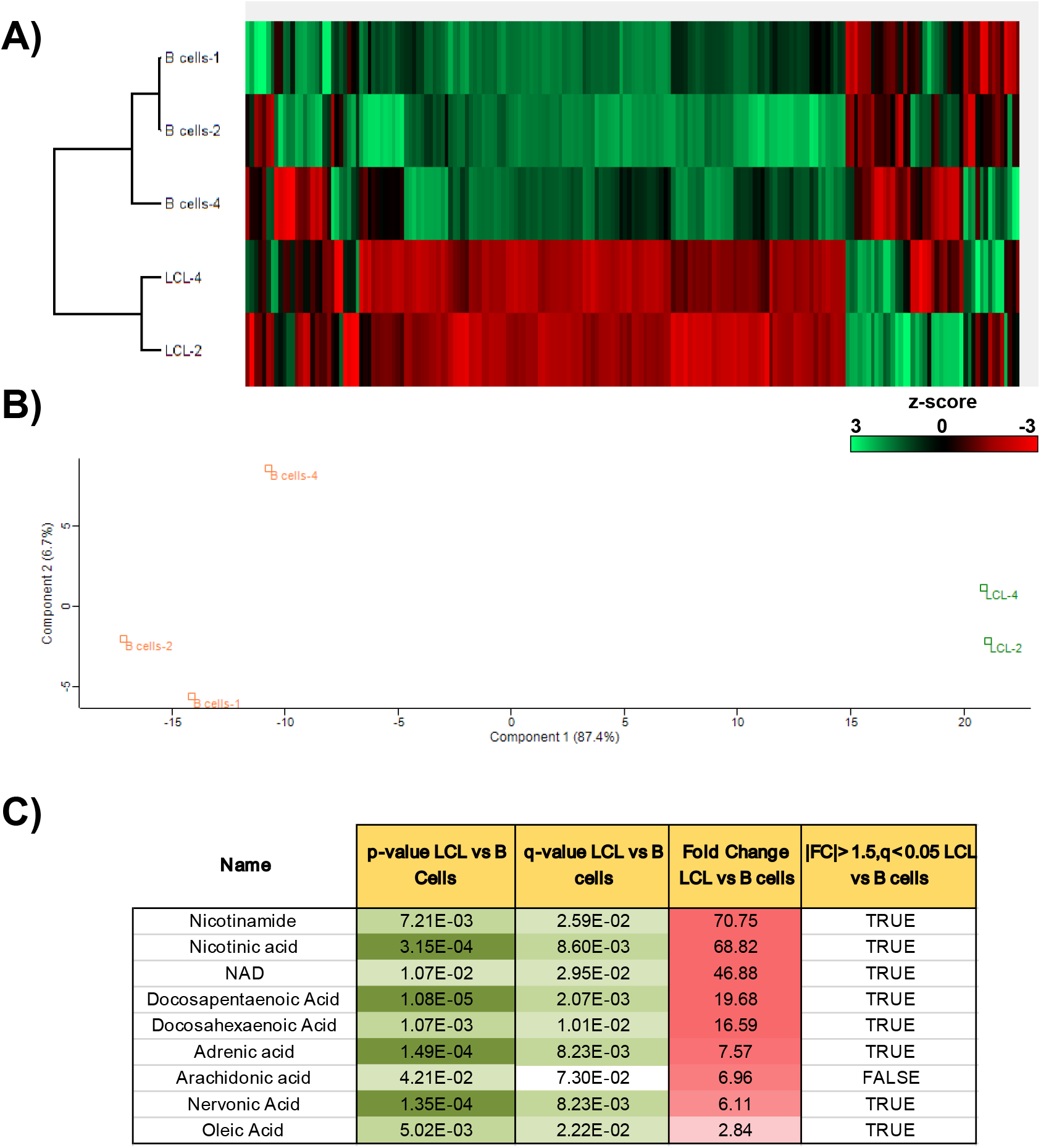
A targeted relative quantitation of approximately 200 polar metabolites spanning 32 different classes examining EBV-immortalization of B cells. **A)** Heat map comparing metabolite levels in primary B cells versus their matched LCLs following EBV-immortalization of B cells 60 days post infection. Heat maps were generated using Perseus software by performing hierarchical clustering on Z-score normalized values using default settings (row and column trees, Euclidean distances, k-means preprocessing with 300 clusters). **B)** Principal component analysis (PCA), performed using default settings on Perseus software, of primary B cells from two donors and three LCLs (two matched to primary B cells) following immortalization of B cells. **C**) Peak areas, representing metabolite levels, were extracted using ThermoScientific Compound Discoverer 3.0. The peak areas were normalized using constant sum. Metabolites were identified from a provided mass list, and by MS/MS fragmentation of each metabolite follow by searching the mzCloud database (www.mzcloud.org). Comparisons between the conditions were performed: Student’s T-test p-value; q-value: Benjamini-Hochberg FDR adjusted p-value to account for multiple testing. q-value < 0.05 is considered significant and flagged with “+” in the “Significant” column; Fold change between 2 conditions (based on average value of the quadruplicate sample); Proteins displaying significant change (q-value < 0.05) with at least 1.5 fold change are indicated in the “FC>1.5, p<p0.05” column.

### EBV-induced immortalization of B cells upregulates FASN and lipogenesis

As we had already determined that LMP1 could induce FASN and lipogenesis in B cells, and both our LMP1 and EBV-immortalization metabolite studies showed significant changes to fatty acids, we also wanted to examine the effect of EBV-induced immortalization of B cells on FASN and lipogenesis. We first extracted proteins from primary B cells and their established LCLs and then assessed FASN protein levels by western blotting analysis. We found was a massive upregulation of FASN at the protein level in LCLs compared to primary B cells (**Fig. 4C**). Specifically, FASN in B cells was barely detectable or not present compared to the robust expression in matched LCLs. Under serum-deprived conditions, we then stained primary B cells and LCLs cells with Nile red followed by FACS analysis (**Fig. 4A**). Similar to our FASN western blot results, we observed virtually no Nile Red staining in B cells and strong staining in our LCLs, which was further confirmed by confocal microscopy imaging (**Fig. 4B**). These findings suggest that EBV-induced immortalization of B cells activates a lipogenesis program as shown by substantial upregulation of fatty acids and their metabolic cofactors, FASN, and lipogenesis. To investigate the dependence of EBV-mediated immortalization on FASN and *de novo* lipid synthesis in B-cells, we performed four independent EBV immortalization assays on primary donor B cells (from three separate donors with information available in **supplementary table 2**), with and without the FASN inhibitor C75 (52). First, 10 million primary B-cells per group were infected with B95.8 strain EBV and left to incubate for 24 hours to allow sufficient time for cell entry and establishment of primary infection. Cells were then treated with 10 μg/mL C75 or equal volume of DMSO and left to incubate for an additional 24 hours, after which time they were imaged via inverted light microscope (**Fig. 4D**). B cells were imaged again at 48 hours post C75 treatment (**Fig. 4E**). Average colony size at 24 hours and 48 hours post C75 treatment was calculated for each pair of four donor B-cells using the “analyze particle” feature of ImageJ software (**Fig. 4D and 4E**). For each control and treated donor set, the average colony size was significantly decreased with FASN inhibition at 24 hours and 48 hours. For both independent immortalization assays of donor 517, B-cell clonal expansion was almost entirely undetectable after 48 hours of FASN inhibition. The average number of colonies per image (30 images per well) was also calculated by setting a size threshold of ≥1000 pixels^2^ as a qualifier of a “healthy, normal cell colony”, as B-cells grow in well-defined “clumps” *in vitro* (**Supplemental Fig 1**). The number of colonies was also significantly higher in the control group than the treatment group for each donor pair. Overall, these results demonstrate that EBV infection induces lipogenesis through FASN, and the inhibition of FASN blocks EBV-induced cell growth transformation of primary B cells.

**Figure 4.**
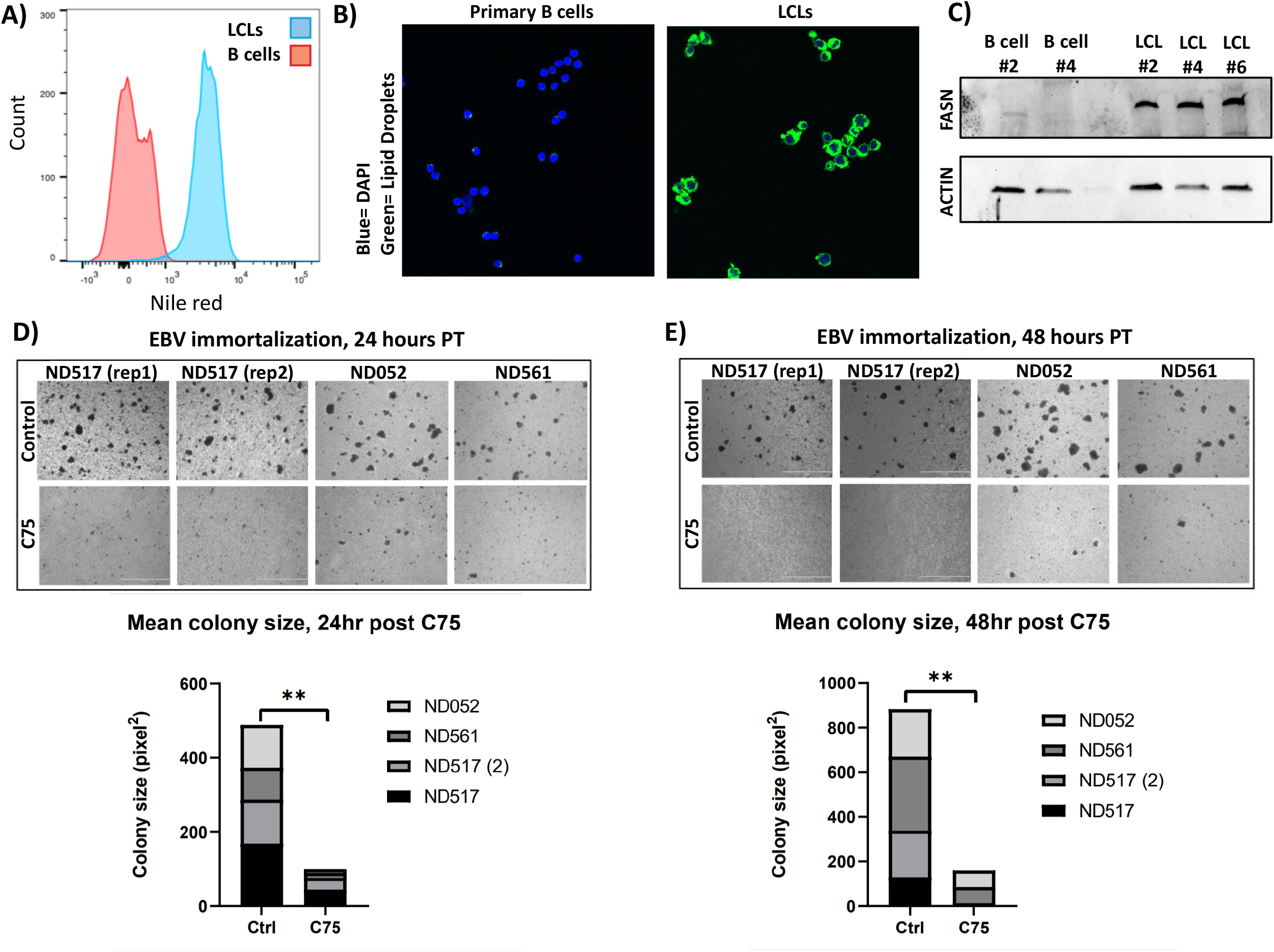
EBV-induced immortalization of B cells upregulates FASN and lipogenesis. **A)** FACs analysis of Nile Red fluorescence staining (excitation, 385 nm; emission,535 nm) for lipid droplets overlaying primary B cells with LCLs. **B)** Confocal microscopy of Nile Red fluorescence staining (excitation, 385 nm; emission,535 nm) for lipid droplets in primary B cells and LCLs. Cells were counterstained with DAPI to stain cell nuclei. **C)** Western blot for FASN in primary B cells and their matched LCLs. Actin served as a loading control. **D-E)** Imaging of primary B-cell EBV immortalization. 10 million cells per group were collected from three donors (one donor was assayed at two independent times) and infected with B95.8 strain EBV 24hours prior to treatment. Cells were imaged on a Nikon TE2000 Inverted Microscope at 4x magnification 24 (D) and 48 hours (E) post C75 treatment. Statistics for average colony size were collected using the “analyze particle” feature of ImageJ for 30 randomized, nonoverlapping images taken of each group. The 30 mean colony size values were then averaged. P values for significant differences (Student’s t-test) are summarized by three asterisks (p<0.001), two asterisks (p<0.01), or one asterisk (p<0.05).

### LMP1+ B cells are more sensitive to FASN inhibition

Dysregulated FASN and lipogenesis is a hallmark of cancer, and cancer cells have been shown to become addicted to the FASN pathway and *de novo* lipogenesis (15). This observation has led to many attempts to target FASN in cancers. Because of this, we sought to examine a potential LMP1-mediated dependency on the FASN pathway by using FASN inhibitors to selectively kill LMP1-expressing cells. Using the FASN inhibitor C75, we generated dose response curves for LMP1-expressing cells vs empty vector control using percent of cell death as determined by a trypan blue exclusion assay. C75 dose concentrations were transformed to log10 prior to nonlinear regression analysis and EC50 values were estimated (**Fig. 5A**). We calculated EC50 values of 72 μM and 36 μM for pBABE and LMP1, respectively, suggesting an increased sensitivity to FASN inhibition in cells expressing LMP1 and increased FASN levels. We then treated latency type I and III cells with FASN inhibitor C75. During various stages of B-cell differentiation *in vivo*, EBV will express either the latency III, II or I program, which entails expression of different subsets of latency genes. Type I latency cells do not endogenously express LMP1 as opposed to latency type III (33, 34). Comparing two such cell types therefore offers a more physiologically relevant comparison between LMP1-positive and negative cells. Mutu I and III are EBV-infected BL cell lines that differ only in their EBV latency status (I vs III). When we treated the LMP1-expressing Mutu III cells with C75, we observed significantly higher cell death compared to Mutu I cells that do not express LMP1 (**Fig. 5B**). Two LCL cell lines (Mutu-LCL and GM12878) also demonstrated sensitivity to FASN inhibition with significant accumulation of cell death after 24 hours compared to DMSO control (**Fig. 5C**). We then measured cell viability following FASN inhibition in primary B-cells and matched LCLs. Whereas uninfected B-cell viability was unaffected by C75 treatment, LCLs showed a significant drop in viability of around 50% vs untreated control (**Fig. 5D**). Cells were also dosed with palmitic acid, which is the predominant product of FASN and was used to determine if the observed toxicity of FASN inhibition was due to lack of fatty acid synthesis or toxic build-up of precursors (35). LCLs responded to palmitic acid with a significant increase in cell viability, given both individually and in combination with C75. This demonstrates that the effects of C75 are due to the halt of downstream fatty acid metabolite synthesis, which is required for the viability of B cells latently infected with EBV.

**Figure 5.**
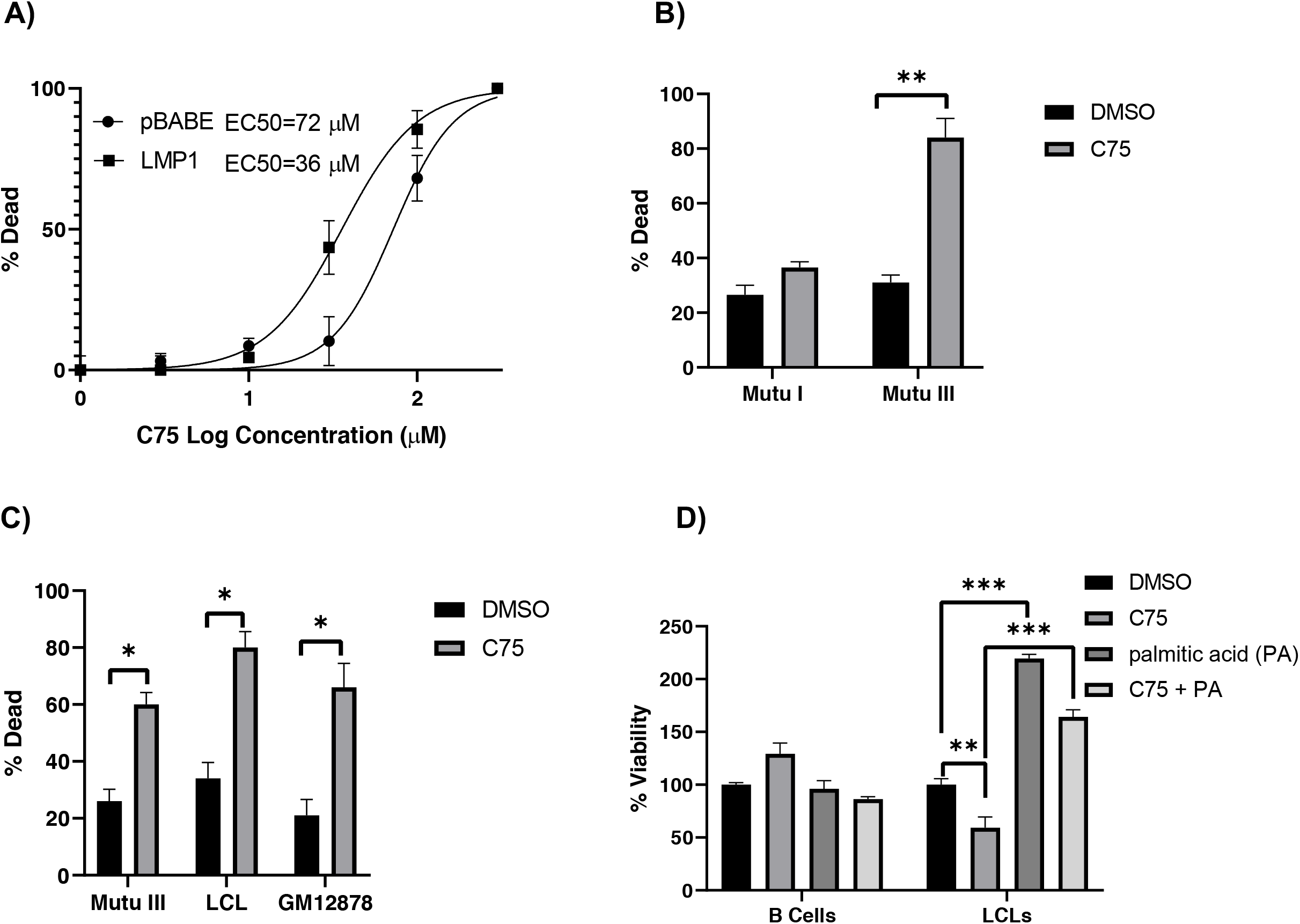
LMP1+ B cells are more sensitive sensitivity to FASN inhibition. **A)** Dose-response curve of DG75 cells that were transduced with retroviral particles containing either pBABE (empty vector) or pBABE-HA-LMP1 vectors and treated with C75 for 24 hrs. Percent of cell death was determined by a trypan blue exclusion assay. Dose concentrations were transformed to log10 prior to nonlinear regression analysis. Data representative of three biological replicates. **B)** Type I (Mutu I) and type III (Mutu III) latently infected EBV-positive B cell lines were incubated with 10 μg/mL of the FASN inhibitor C75 or DMSO control for 24 hrs. Percent of cell death as determined by a trypan blue exclusion assay. **C)** Type III latently infected EBV-positive B cell lines were incubated with 10 μg/mL of the FASN inhibitor or DMSO control for 24 hrs. Percent of cell death as determined by a trypan blue exclusion assay. **D)** Primary B cells and LCLs were incubated with 10 μg/mL of the FASN inhibitor C75, 25 μM palmitic acid (PA), C75+PA or DMSO control for 24 hrs. Cell viability was determined by cell titer glo assay. Error bars represent standard deviation of two independent experiments. P values for significant differences (Student’s t-test) are summarized by three asterisks (p<0.001), two asterisks (p<0.01), or one asterisk (p<0.05).

### LMP1 stabilizes FASN protein levels

We next sought to determine the mechanisms LMP1 employs to upregulate FASN. Previous work has pointed to LMP1 driving expression of FASN through its upstream regulator SREBP1c, at least in the context of NPC (27). However, our previously published RNA-seq data (9) did not suggest that SREBP1c was a factor upregulated by LMP1, and this was confirmed by RT-qPCR using primers against both the precursor and mature isoforms of SREBF, SREBFa and SREBFc (**Fig. 6B**). In fact, both FASN and SREBFa were downregulated in LMP1+ cells vs LMP1-cells while SREBFc remained unchanged. However, FASN can be stabilized at the protein level by USP2a, a ubiquitin-specific protease that functions by removing ubiquitin from FASN and thus prevents its degradation by the proteasome (26) (**Fig. 6A**). Our RNA-seq dataset (9) suggested that USP2a is upregulated by LMP1 and this was confirmed by RT-qPCR (**Fig. 6B**). Because of this, we then wanted to determine if LMP1 stabilized FASN at the protein level. To examine the effect of LMP1 expression of FASN protein levels, we used the protein synthesis inhibitor cycloheximide (CHX). Following treatment with CHX, we observed that FASN protein levels were more stable at 24 hours in our LMP1-expresssing cell line vs empty vector control (**Fig. 6D**). This suggests that ectopic expression of LMP1 can induce the post-translational stabilization FASN in BL cell lines. To examine potential mechanisms of how EBV infection was causing upregulation of FASN, we again looked at factors effecting both expression and post-translational modifications of the enzyme. We investigated the SREBPs, the principal upstream regulators of FASN gene expression, and USP2a, the ubiquitin-specific protease that stabilizes FASN protein by decreasing its ubiquitination. First, we used RT-qPCR to examine the gene expression of FASN and USP2a. When we compared the expression of these genes between a limited set of matched primary B cells and LCLs, we found interesting results. Depending on the LCL (each generated from a different donor’s B cells) we found that either FASN expression was increased or USP2 expression, but never the two together (**Fig. 6C**). Again, all LCLs robustly upregulated FASN at the protein level, suggesting that EBV will co-opt alternative pathways to achieve the same result of increased FASN abundance. To further investigate FASN protein stability, we performed an immunoprecipitation of FASN and immunoblotted for USP2a (**Fig. 6E**). In both DG75 cells with empty vector or ectopically expressing LMP1, FASN co-immunoprecipitated with USP2a. These results indicate that USP2a stabilizes FASN levels in both cell types, but significantly more-so in those expressing LMP1, even when normalized to their higher basal FASN levels (**Supplemental figure 2**). Furthermore, when LMP1-positive DG75 cells were treated with the USP2a inhibitor ML364, FASN levels are decreased in a dosage-dependent manner (**Fig. 6F** and **Fig. 6G**). Interestingly, in DG75 cells not ectopically expressing LMP1, a rebound effect of FASN levels is observed when ML364 dose is increased from 10μM to 20μM. Finally, when treated with ML364, LMP1-expressing DG75 cells were significantly more sensitive to USP2a inhibition than those with the empty expression vector (**Fig. 6H**). While empty-vector DG75 proliferation rate was decreased ~30% compared to DMSO control at both 10μM and 20μM ML364, LMP1 expressing DG75 proliferation rates were decreased ~70-75%, respectively (**Supplemental figure 3**). Taken together, these results indicate that LMP1-expressing DG75 cells rely on the USP2a mediated post-translational stabilization of FASN protein.

**Figure 6.**
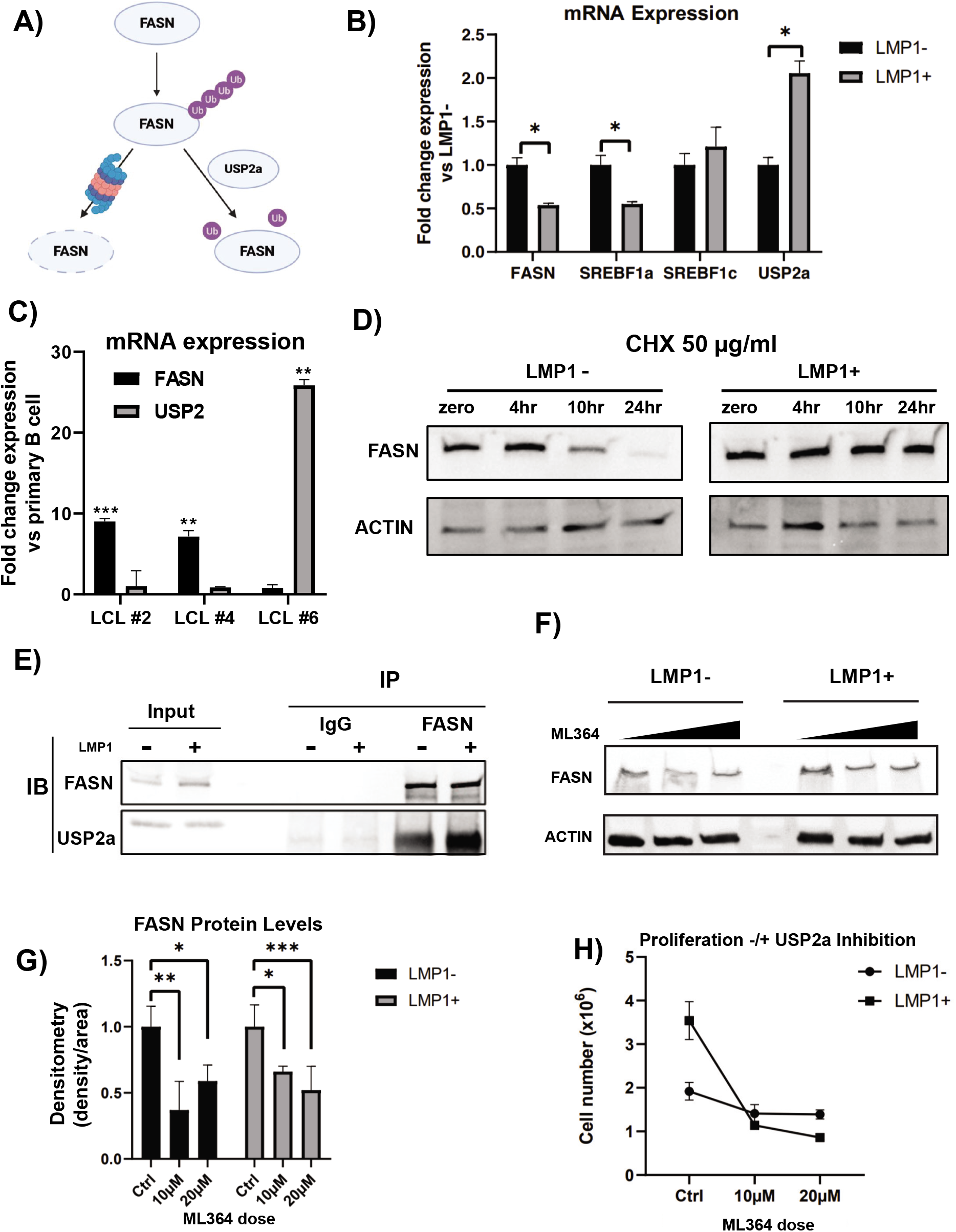
LMP1 stabilizes FASN protein. **A)** Schematic of FASN stabilization. USP2a, a ubiquitin-specific protease, functions by removing ubiquitin from FASN and thus prevents its degradation by the proteasome. **B)** Relative mRNA expression in LMP1 + cells versus empty vector (pBABE) as determined by RT-qPCR using double delta Ct analysis and normalized to 18s. **C)** Relative mRNA expression in LCLs versus primary B cells (3 independent donors) as determined by RT-qPCR using double delta Ct analysis and normalized to 18s. **D)** FAS protein levels in LMP1- and LMP1+ cells treated with 50 μg/mL cycloheximide over a 24-hour time course. Actin was included as a loading control. **E)** 10 μg of polyclonal rabbit antibody to FASN or normal rabbit IgG was added to the lysate of 10 million LMP1- or LMP1+ DG75 cells, respectively. Magnetic protein-A conjugated beads were utilized to immunoprecipitate FASN/IgG binding proteins. Beads were boiled in 2x laemmli buffer and ran on a western blot beside 10% protein lysate input and blotted for FASN and USP2a signal. **F)** 5 million LMP1- and LMP1+ DG75 cells were treated with DMSO control (0μM) or the USP2a inhibitor ML364 at 10μM or 20μM. Cell lysates were collected after 24 hours and ran on a western blot. Blots were probed for FASN and actin as a control. Graph is representative of signal density/area of FASN, normalized to actin control. DMSO control was set to 1, with treatment groups displayed as fold change relative to DMSO control. G) 1 million LMP1- or LMP1+ DG75 cells were treated with DMSO control, 10μM ML364, or 20μM ML364. At 24 hours, cells were counted with trypan blue to exclude dead cells. Statistics of each treatment group are comparing the difference between proliferation rates with regard to LMP1 expression. Error bars represent standard deviation of three independent experiments. P values for significant differences (Student’s t-test) are summarized by four asterisks (p≤ 0.0001), three asterisks (p<0.001), two asterisks (p<0.01), or one asterisk (p<0.05).

## Discussion

The EBV-encoded oncoprotein LMP1 is expressed in several EBV associated malignancies, including Hodgkin and post-transplant B-cell lymphomas and NPC. We and others have previously reported that LMP1 can stimulate aerobic glycolysis (‘Warburg’ effect) in cells (8–13). Our initial work was grounded in expression data, where we observed that LMP1 could induce HIF-1α-dependent gene expression, alteration of cellular metabolism, and accelerated cellular proliferation (9). As a follow up to further investigate these LMP1-associated cellular metabolic changes, we used a targeted approach to examine the effects of both the ectopic expression of LMP1, as well as EBV-mediated B-cell growth transformation, on host metabolites. We observed that the top 15-20 metabolites significantly induced by LMP1 in the BL cell line DG75 were fatty acids from this initial analysis. The observed induction of fatty acids aligned with increased levels of FASN and lipid droplet formation as compared to empty vector controls. A recent study has specifically linked LMP1 to the promotion of *de novo* lipogenesis, lipid droplet formation, and increased FASN in NPC (27). This study went on to show that FASN overexpression is common in NPC, with high levels correlating significantly with LMP1 expression. Moreover, elevated FASN expression was associated with aggressive disease and poor survival in NPC patients. Interestingly, alteration of lipid metabolism was also observed in Burkitt Lymphoma following gene expression analysis. Based on this, adipophilin was identified as a novel marker of BL (28). This elevated level of lipid metabolism in BL might explain why we observed relatively minor changes to FASN levels and lipid droplet formation when we introduced LMP1 to the EBV-negative BL cell line DG75.

Additionally, the increase in fatty acids via ectopic expression of LMP1 was offset following treatment with the PARP inhibitor olaparib. Previously, we have shown that PARP1 is important in LMP1-induced aerobic glycolysis and accelerated cellular proliferation, both of which could be attenuated with PARP inhibition. PARP1 gene deletion and inhibition have been reported to enhance lipid accumulation in the liver and exacerbate high fat-induced obesity in mice (36, 37). However, a conflicting report concludes robust increases in PARP activity in livers of obese mice and non-alcoholic fatty liver disease (NAFLD) patients and that inhibition of PARP1 activation alleviates lipid accumulation and inflammation in fatty liver (38). Therefore, the role of PARP1 in lipid metabolism remains inconclusive, at least in the context of the liver and diet-induced obesity. As we have previously demonstrated, PARP1 can act as a coactivator of HIF-1α-dependent gene expression. It is of interest to note that an emerging body of work shows that HIF-1α can regulate lipid metabolism (39) including an ability to regulate FASN (40). It still needs to be elucidated, however, how much of the LMP1-mediated changes to aerobic glycolysis and lipid metabolism is facilitated distinctly through PARP1, HIF-1α, a combination of the two, or completely independent of these factors.

In addition to examining LMP1-specific metabolic effects, we then examined metabolic changes following EBV-mediated B-cell growth transformation. While we did not find that all the absolute highest fold changes in metabolites were fatty acids as we did with ectopic expression of LMP1, we did find fatty acids being amongst the top metabolites altered. A recent study used proteomics to examine resting B-cells and several time points after EBV infection. Their data pointed to the induction of one-carbon (1C) metabolism being necessary for the EBV-mediated B-cell growth transformation process (41). This same analysis also revealed that EBV significantly upregulates fatty acid and cholesterol synthesis pathways. There are several key differences in this proteomics study versus our metabolomics approach. While we compared resting B-cells with established LCLs around two months after infection, the above study also used several earlier timepoints. A follow-up study using the same dataset suggested essential roles for Epstein-Barr nuclear antigen 2 (EBNA2), SREBP, and MYC in cholesterol and fatty acid pathways (42). The EBV-encoded transcription factor EBNA2 is produced early in the infection phase (72hrs) (43), and the cholesterol and fatty acids synthesis pathways, including upregulation of FASN, were found to be induced early in infection (96 hours). As LMP1 appears after 3-7 days post-infection (43), the role of LMP1 in the induction of the referenced pathways remains unclear. The study mentioned above (42) indicated an important role for Rab13 role in the possible trafficking of LMP1 to lipid raft signaling sites. Therefore, it is possible that the early changes to cholesterol and fatty acids synthesis pathways aid in the localization of LMP1 to cellular membranes, enabling LMP1 to maintain cholesterogenic and lipogenic programs at later timepoints by stimulating PI3K/AKT signaling cascades. These studies also provide rationale for our EBV-immortalization assays of primary donor B-cells with and without FASN inhibition at 48 hours post-infection. This timepoint would precede the induction of both FASN and LMP1. By inhibiting FASN, and thus *de novo* lipogenesis before LCL-associated addiction to various metabolic pathways can be established, we can conclude its essential role in this process.

Outside of the context of EBV-mediated B-cell growth transformation, there is also evidence of glucose-dependent de novo lipogenesis in B-lymphocytes following lipopolysaccharide (LPS)-stimulated differentiation into Ig-secreting plasma cells (44). Specifically, this study pointed to ATP citrate lyase (ACLY) linking glucose metabolism into fatty acid and cholesterol synthesis during differentiation. This becomes especially interesting when considering the ability of EBV and LMP1 to induce both aerobic glycolysis and lipogenesis programs. One of the questions that arise from such studies is: are these metabolic changes unique to EBV-induced immortalization of B-cells, or are we observing the hijacking of pathways and metabolic remodeling used in the normal proliferation and differentiation of B-cells? A study into primary effusion lymphoma (PEL) cells, which are a unique subset of human B-cell non-Hodgkin lymphomas cells latently infected with Kaposi’s sarcoma-associated herpesvirus (KSHV, another γ-herpesvirus), showed that FASN expression and induction of fatty acid synthesis was necessary for the survival of latently infected PEL cells (45). Interestingly and related to the aforementioned question, these researchers stimulated resting B-cells with LPS to determine if differences in glycolysis and FASN were a consequence of proliferation, as PEL cells are continuously proliferating as lymphomas, rather than the transformed phenotype. While they did observe an elevated rate of glycolysis following LPS-stimulation of primary B-cells, it was still significantly lower than that of vehicle-treated PEL cells. In addition, FASN did not substantially change in LPS-stimulated versus vehicle-treated primary B-cells; nor did LPS stimulation of PEL lead to any further increases in glycolysis or FASN compared with vehicle-treated PEL (45). These data potentially suggest that FASN activity is an independent phenotype of γ-herpesvirus, whether in the context of latently infected KSHV PEL or latently infected EBV NPCs and lymphomas, rather than a consequence of increased proliferation index.

We then went on to show that LMP1-expressing cells, including those ectopically expressing LMP1, latency type III cell lines, and LCLs transformed from primary B cells, were all more sensitive to FASN inhibition vs their corresponding LMP1-negative controls. Analysis of FASN expression in NPC patients found that higher levels of FASN expression significantly correlated with advanced primary tumor and distant lymph node metastasis (27). Latent infection of endothelial cells by KSHV led to a significant increase in long-chain fatty acids as detected by a metabolic analysis. Fatty acid synthesis is required for the survival of latently infected endothelial cells, as inhibition of key enzymes in this pathway led to apoptosis of infected cells (46). We also observed that primary B-cells, which express no or very little FASN protein, unsurprisingly were not sensitive to FASN inhibition. However, our LCLs transformed from primary B-cells developed sensitivity to FASN inhibitors corresponding to FASN and lipogenesis induction. We also showed that FASN inhibition via C75 ablated the ability of EBV to immortalize primary B-cells. A study reported that the use of the lipoprotein lipase inhibitor orlistat resulted in apoptosis of B-cell chronic lymphocytic leukemia (CLL) cells without killing normal B-cells from donors (47).

Finally, we observed somewhat surprising results when one donor LCL displayed hugely upregulated USP2a mRNA compared to its matched primary B-cell as well as the other matched LCL/B-cell pairs. Conversely, donor LCL #6 had relatively lower FASN mRNA levels compared to the other donor LCLs. Considering that FASN levels can be regulated both transcriptionally and post-translationally, we sought to investigate the mechanism different LCLs employ to maintain relatively high FASN protein levels. First, we showed that FASN and USP2a bind in human B-cells, utilizing LMP1 or empty vector DG75 BL cells. We found that not only do USP2a and FASN interact in both lines, but stronger/more frequently in LMP1-expressing cells. This suggests to us that while the relationship between the proteins is not entirely dependent on EBV, it is strengthened by LMP1. Utilizing the same two cell lines, we also showed that inhibition of USP2a via the drug ML364 significantly decreased FASN protein levels in a dose-dependent manner in the LMP1-positive DG75. While FASN levels were also decreased in the empty vector DG75, there was a slight rebound effect observed when ML364 dosage was increased from 10 μM to 20μM. This again indicates that LMP1 selectively employs USP2a to stabilize FASN. Finally, while the proliferation of empty vector DG75 decreased roughly 30% at both ML364 concentrations, LMP1-positive DGs experienced a 70-75% decrease, respectively. Not only is LMP1+ proliferation significantly decreased by USP2a inhibition, but it is also considerably reduced compared to empty vector DG75 cells at the same dosage. From this, we can conclude that USP2a inhibition selectively inhibits the proliferation of LMP1-positive BL cell lines, providing rationale into a future investigation of ML364 treatment of LMP1-positive malignancies, and solidifying another example of USP2a-induced stabilization of FASN in a third, separate human cancer.

In conclusion, LMP1 is expressed in most EBV-positive lymphomas, and EBV-associated malignancies are often associated with a worse prognosis than their EBV-negative counterparts. Despite many attempts to develop novel therapies, EBV-specific treatments currently remain largely investigational. Therefore, there is an apparent demand for EBV-specific therapies for both prevention and treatment. The work presented here suggests that targeting lipogenesis programs may be an effective strategy in the treatment of LMP1-positive EBV-associated malignancies. Further studies into the metabolic signaling pathways manipulated by EBV is critical to aid in the development of targeted, novel therapies against EBV-associated malignancies.

## Materials and Methods

### Cell culture and drug treatment

All cells were maintained at 37 °C in a humidified 5% CO2 atmosphere in medium supplemented with 1% penicillin/streptomycin antibiotics. Lymphocyte cell lines (EBV-negative Burkitt’s lymphoma cell line DG75 ATCC CRL-2625 (DG75), EBV-positive latency III cell lines Mutu III, Mutu-LCL, GM12878 and EBV-positive latency I cell line Mutu I) were cultured in suspension in RPMI 1640 supplemented with fetal bovine serum at a concentration of 15%. Primary B cells were cultured in suspension in RPMI 1640 supplemented with fetal bovine serum at a concentration of 20%. 293T ATCC CRL-3216 (HEK 293T) cells were cultured in Dulbecco’s modified Eagle medium (DMEM) supplemented with fetal bovine serum at a concentration of 10%. Olaparib 5μM (Selleck Chemical), cycloheximide 50 μg/mL (Sigma), C75 10μg/mL (Sigma), and ML364 10μm/20μM (selleckchem) was dissolved in dimethyl sulfoxide (DMSO) when used in respective in vitro assays.

### Retroviral transduction

Plasmid constructs hemagglutinin (HA)-tagged full-length LMP1, pBABE, pVSV-G, and pGag/Pol were kindly provided by Nancy Raab-Traub (UNC, Chapel Hill, NC) and were described previously [59]. Retroviral particles were generated using the Fugene 6 reagent (Promega) to simultaneously transfect subconfluent monolayers of 293T cells with 1μg pBABE (vector) or HA-LMP1, 250 ng pVSV-G, and 750 ng pGal/Pol according to the manufacturer’s instructions. Supernatant containing lentivirus was collected at 48- and 72-h post-transfection and filtered through a 0.45 μM filter. DG75 cells were transduced by seeding 5×105 cells in 6-well plates in 500 μl medium and adding 500 μl of medium containing retroviral particles. The transduced cells were placed under long-term selection in medium containing 1 μg/ml puromycin.

### EBV Infection of Primary B cells

De-identified, purified human B cells were obtained from the Human Immunology Core of the University of Pennsylvania under an Institutional Review Board-approved protocol and were isolated using the RosetteSep Human B Cell Enrichment Cocktail (StemCell Technologies) as per protocol. Primary B cells were infected with concentrated B95.8 strain EBV within 24 hours of their purification from donor plasma. EBV was collected from supernatant of the EBV-positive ATTC cell line VR-1492TM, which was concentrated with PEG Virus Precipitation Kit (abcam). Infected cells were for cultured for 60 days before being considered a lymphoblastoid cell line for all assays compared against matched primary B-cells. In evaluating the role of FASN in EBV immortalization, B-cells were infected with concentrated EBV for 24 hours before being treated with 10μg/mL of C75 or equal volume DMSO.

### Targeted relative metabolites quantitation

Cells were pelleted by centrifugation at 2,000 rpm for 5 min, 4 °C, washed cells twice in ice-cold PBS. Samples were extracted using cold extraction solution containing 80% methanol/20% water/0.2 uM heavy internal standard mix (MSK-A2-1.2 Cambridge Isotope Laboratories, Inc) using 2 million cells in 500 uL. Samples were vortexed thoroughly for 30 sec and placed on dry ice for at least 15 min. Samples were then spun at max speed (>13,000 rpm) for 15 min at 4 °C to pellet any debris. LC-MS analysis was performed on a Thermo Fisher Scientific Q Exactive HF-X mass spectrometer equipped with a HESI II probe and coupled to a Thermo Fisher Scientific Vanquish Horizon UHPLC system. Polar metabolites were extracted using 80% methanol and separated at 0.2 ml/min by HILIC chromatography at 45 °C on a ZIC-pHILIC 2.1 inner diameter x 150-mm column using 20 mM ammonium carbonate, 0.1% ammonium hydroxide, pH 9.2, and acetonitrile with a gradient of 0 min, 85% B; 2 min, 85% B; 17 min, 20% B; 17.1 min, 85% B; and 26 min, 85% B. Relevant MS parameters were as follows: sheath gas, 40; auxiliary gas, 10; sweep gas, 1; auxiliary gas heater temperature, 350 °C; spray voltage, 3.5 kV for the positive mode and 3.2 kV for the negative mode; capillary temperature, 325 °C; and funnel RF level at 40. A sample pool (quality control) was generated by combining an equal volume of each sample and analyzed using a full MS scan at the start, middle, and end of the run sequence. For full MS analyses, data were acquired with polarity switching at: scan range 65 to 975 m/z; 120,000 resolution; automated gain control (AGC) target of 1E6; and maximum injection time (IT) of 100 ms. Data-dependent MS/MS was performed without polarity switching; a full MS scan was acquired as described above, followed by MS/MS of the 10 most abundant ions at 15,000 resolution, AGC target of 5E4, maximum IT of 50 ms, isolation width of 1.0 m/z, and stepped collision energy of 20, 40, and 60. Metabolite identification and quantitation were performed using Compound Discoverer 3.0. Metabolites were identified from a mass list of 206 verified compounds (high confidence identifications) as well as by searching the MS/MS data against the mzCloud database and accepting tentative identifications with a minimum score of 50.

### Lipid droplet fluorescence staining

Nile Red fluorescence staining was assessed with the Lipid Droplets Fluorescence Assay Kit according to the manufacturer’s protocol (Cayman Chemical, Ann Arbor, MI, USA). One day before staining assay, cells were incubated in serum free medium. As a positive control, cells in completed medium were treated overnight with Oleic Acid provided from assay kit at 1:2000 dilution. For lipid droplets staining and quantification using a plate reader, cells were fixed with 1X assay fixative, washed with PBS and then stained with working solution of Hoechst 33342 (1 ug/ml)) and Nile Red (1:1000). The fluorescence of cells was determined using a GloMax plate reader (Promega). Hoechst 33342 fluorescence was measured with an excitation of 355 nm and an emission of 460 nm, while Nile Red fluorescence was determined using a 485 nm excitation and 535 nm emission. Differences in cell number were corrected by using Hoechst 33342 fluorescence signal to normalize the Nile Red signal in each well. For flow cytometric analysis, cells were only stained with Nile Red (1:1000). Analysis was carried out using a FACS Calibur flow cytometer (Becton Dickinson) and CellQuest software, and the cell population was analyzed using FlowJo software. Confocal microscopy images were taken on a Leica TCS SP8 MP multiphoton microscope.

### Western Blot Analysis

Cell lysates were prepared in radioimmunoprecipitation assay (RIPA) buffer (50 mM Tris-HCl, pH 7.4, 150 mM NaCl, 0.25% deoxycholic acid, 1% NP-40, 1 mM EDTA) supplemented with 1X protease inhibitor cocktail (Thermo Scientific). Protein extracts were obtained by centrifugation at 3,000×g for 10 minutes at 4 °C. For nuclear fractionation, nuclear soluble and chromatin-bound protein fractions were extracted from cells using the Subcellular Protein Fractionation Kit for Cultured Cells kit (Invitrogen) according to manufacturer’s instructions. The bicinchoninic (BCA) protein assay (Pierce) was used to determine protein concentration. Lysates were boiled in 2x SDS-PAGE sample buffer containing 2.5% β-mercaptoethanol, resolved on a 4 to 20% polyacrylamide gradient Mini-Protean TGX precast gel (Bio-Rad), and transferred to an Immobilon-P membrane (Millipore). Membranes were blocked for 1 h at room temperature and incubated overnight with primary antibodies recognizing LMP1 (Abcam ab78113), FASN (Abcam ab22759), USP2a (Abcam ab66556), and Actin (Sigma A2066), as recommended per the manufacturer. Membranes were washed, incubated for 1 h with the appropriate secondary antibody, either goat anti-rabbit IgG-HRP (Santa Cruz sc-2030) or rabbit anti-mouse IgG-HRP (Thermo Scientific 31430). Membranes were then washed and detected by enhanced chemiluminescence.

### Co-immunoprecipitation

For FASN-immunoprecipitation (IP) assays, 10 million empty vector or pBabe-LMP1 DG75 cells were collected for each IP and resuspended in 1mL of RIPA buffer with protease/phosphatase inhibitor cocktail (Thermo Scientific). Before addition of 10ug of either FASN (abcam, ab99359) or normal rabbit IgG (Jackson, 111-005-003), 50uL of cell lysate was collected and kept as input material. Cell lysates were incubated with respective antibodies for one hour at room temperature, rotating, after which 30uL of protein A magnetic beads (Invitrogen, 10001D) were added. The mixture was left to incubate overnight at 4°, rotating. The beads were then separated with a magnetic rack and washed three times in RIPA buffer with protease/phosphatase inhibitor, each for 10 minutes in a 4° thermomixer at 1000rpm. The beads were then boiled at 95° for 8 minutes in 50uL 2x laemmli buffer, with half of the volume ran on an immunoblot for FASN, and half for USP2a (abcam, ab66556) as described above. Densitometry analysis was performed on Invitrogen iBright Analysis Software, with signal density/area from IgG control lanes subtracted from IP lanes. IgG normalized IP signal was then normalized to input signal density/area. Data shown is representative of three independent co-IP assays, averaged.

### RT-qPCR

For reverse transcription quantitative PCR (RT-qPCR), RNA was extracted from 2 × 106 cells using TRIzol (Thermo Fisher Scientific) according to the manufacturer’s instructions. SuperScript II reverse transcriptase (Invitrogen) was used to generate randomly primed cDNA from 1 μg of total RNA. A 50-ng cDNA sample was analyzed in triplicate by quantitative PCR using the ABI StepOnePlus system, with a master mix containing 1X Maxima SYBR Green and 0.25 μM primers.

Data were analyzed by the ΔΔCT method relative 18s and normalized to untreated controls. Primers are available upon request.

### Cell Viability Assay

Cell viability was measured using the CellTiter-Glo Luminescent Cell Viability Assay (Promega). 100 μl of cells in culture medium per well were plated in 96-well opaque-walled plates. The plate and samples were equilibrated by placing at room temperature for approximately 30 minutes. 100 μl of CellTiter-Glo Reagent was added to 100μl of medium containing cells. Plate contents for were then mixed for 2 minutes on an orbital shaker to induce cell lysis. Finally, the plate was incubated at room temperature for 10 minutes to stabilize luminescent signal before luminescence was recorded on a GloMax plate reader (Promega).

### Dose-response curves

Dose concentrations were transformed to log10 prior to nonlinear regression analysis using GraphPad Prism version 8.00 for Mac OS X, GraphPad Software, La Jolla California USA, www.graphpad.com. Specifically, % dead cells based on live/dead counting using a Countess II FL Automated Cell Counter (ThermoFisher) following incubation with trypan blue was used as the Y value response.

## Acknowledgements

Research reported in this publication was supported by the National Institute of Allergy and Infectious Diseases of the National Institutes of Health under Award Number R01AI130209. The metabolomics analysis was performed at The Wistar Institute Proteomics and Metabolomics Shared Resource on a Thermo Q-Exactive HF-X mass spectrometer purchased with NIH grant S10 OD023586.

