## Supplemental Figures and Table for "Epstein–Barr virus-encoded latent membrane protein 1 and B-cell growth transformation induces lipogenesis through fatty acid synthase"

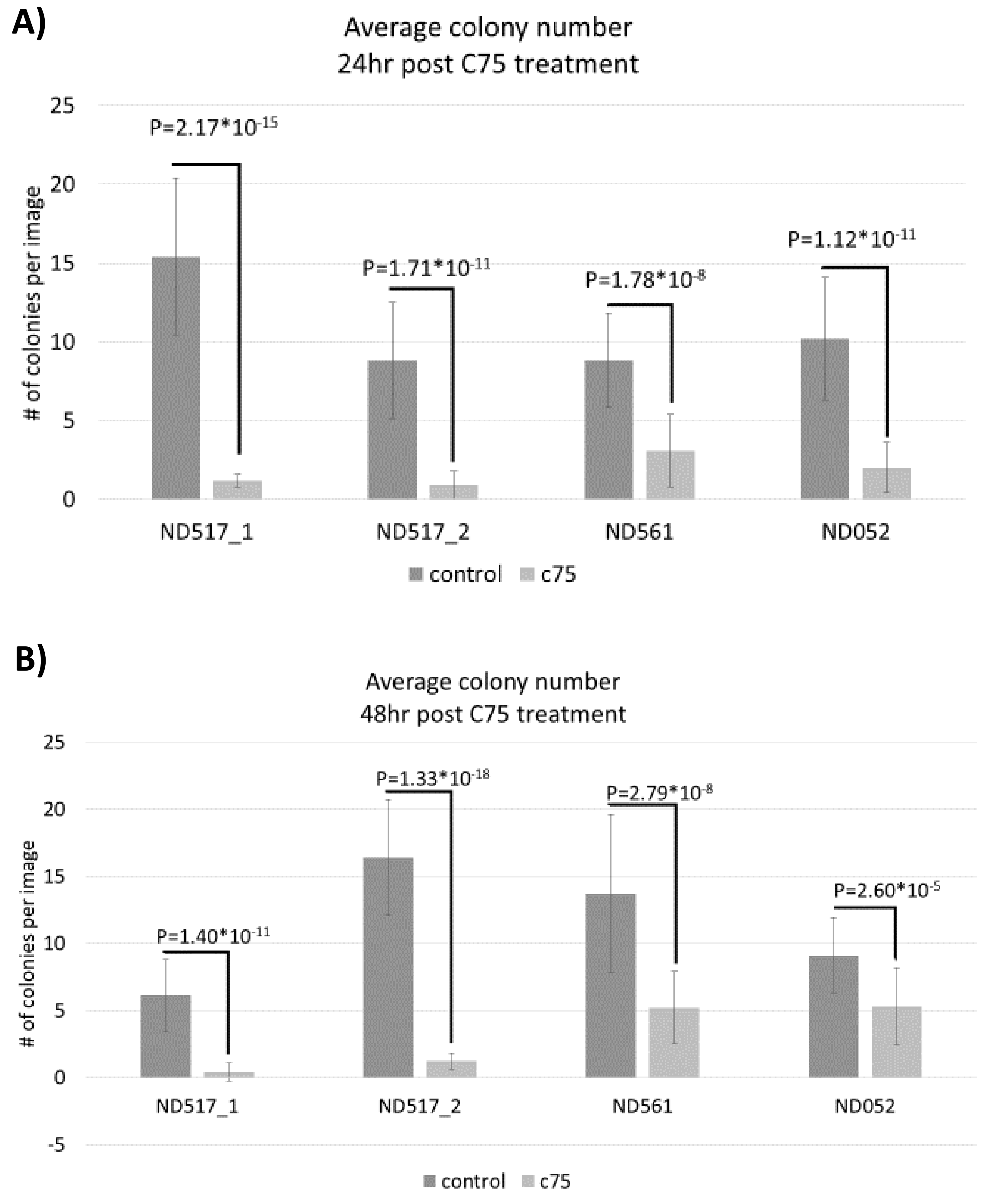
A)
Average colony number
24hr post C75 treatment
### of colonies per image
25
20
15
10
5
0
P=2.17*10-15
P=1.71*10-11
P=1.78*10-8
P=1.12*10-11
ND517_1
ND517_2
ND561
ND052
control
c75
B)
Average colony number
48hr post C75 treatment
### of colonies per image
25
20
15
10
5
0
-5
P=1.40*10-11
P=1.33*10-18
P=2.79*10-8
P=2.60*10-5
ND517_1
ND517_2
ND561
ND052
control
c75

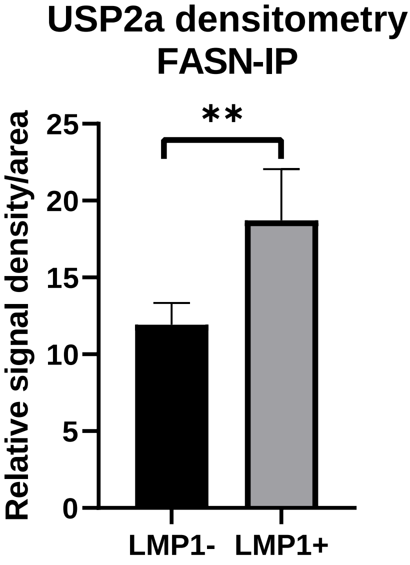
USP2a densitometry
FASN-IP
**
25
20
15
10
5
0
Relative signal density/area
LMP1-
LMP1+

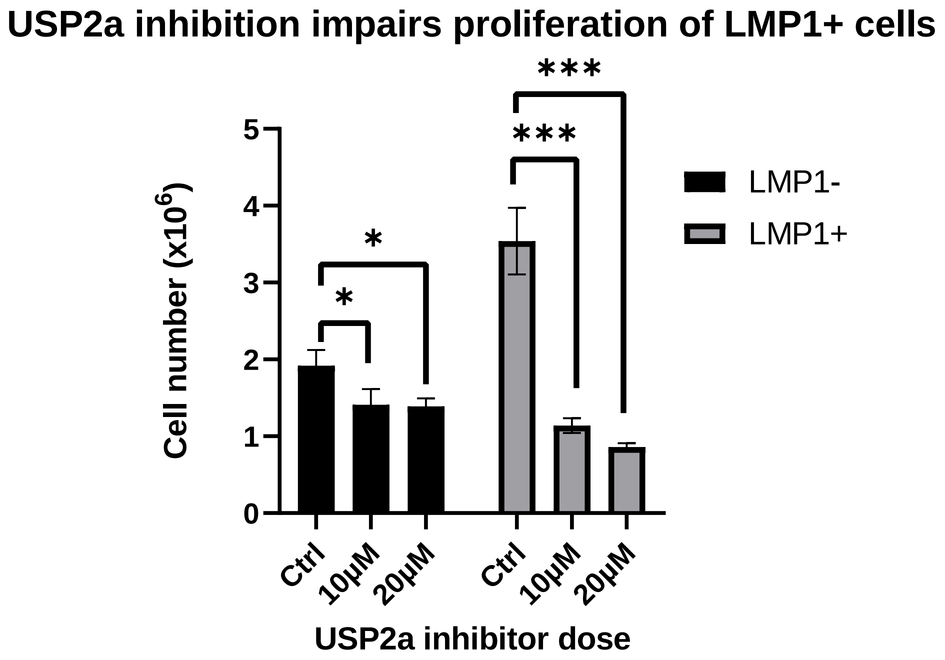

USP2a inhibition impairs proliferation of LMP1+ cells
Cell number (x10^6)
0
1
2
3
4
5
*
*
***
***
LMP1-
LMP1+
Ctrl
10μM
20μM
Ctrl
10μM
20μM
USP2a inhibitor dose

| Donor number | Age | Sex | HLA-A | HLA-B | HLA-C |
|---|---|---|---|---|---|
| ND500 (B-cells 1) | 25 | F | A 02:06, 11:01 | B 15:02, 40:01 | C 04:03, 08:01 |
| ND560 (B-cells 2) | 21 | F | A 02:06, 24:02 | B 15:25, 40:02 | C 03:04, 04:03 |
| ND492 (B-cells 4) | 37 | M | A 01:01, 03:01 | B 07:02, 37:01 | C 06:02, 07:02 |
| ND451 (B-cells 6) | 31 | F | A 29:01, 68:01 | B 15:17, 39:01 | C 07:01, 12:03 |
| ND052 | 56 | M | A2 | B62, B44 | N/A |
| ND517 | 28 | F | A 11: 01, 30:02 | B 18: 01, B 51:01 | C 05: 01, B 15:02 |
| ND561 | 25 | M | A 23:01, 30:01 | B 13:02, 49:01 | C 06:02, 07:01 |

**Supplemental table 1. Complete metabolite comparison of primary B-cells vs matched LCLs.**

**Supplemental table 2. Donor information for primary B-cells.**

De-identified, purified human B-cells were obtained from the Human Immunology Core of the University of Pennsylvania, where HLA typing was performed. Age, sex, and MHCI types are provided above to show heterogeneity of donors.

**Supplemental Figure 1. Cell colony size is decreased by FASN inhibition during EBV-immortalization.** 10 million cells per group were collected from three donors (one donor was assayed at two independent times) and infected with B95.8 strain EBV 24hours prior to treatment. Cells were imaged on a Nikon TE2000 Inverted Microscope at 4x magnification 24 (A) and 48 hours (B) post C75 treatment. 30 randomized, non-overlapping images from each group were collected and analyzed on ImageJ. Considering the average colony size of a healthy cell formation, a threshold of ≥1000 pixels$^2$ was set before using the "analyze colony" feature to determine number of cell formations per image that met the "healthy threshold". The values from the 30 images were then averaged to determine the number of colonies formed on average per donor group per image.

**Supplemental Figure 2. USP2a stabilizes FASN protein in DG75 cells.** 10 μg of polyclonal rabbit antibody to FASN or normal rabbit IgG was added to the lysate of 10 million LMP1- or LMP1+ DG75 cells, respectively. Magnetic protein-A conjugated beads were utilized to immunoprecipitate FASN/IgG binding proteins. Beads were boiled in 2x laemmli buffer and ran on a western blot beside 10% protein lysate input and blotted for FASN and USP2a signal. Densitometry analysis was performed on Invitrogen iBright Analysis Software, with signal density/area from IgG control lanes subtracted from IP lanes. IgG normalized IP signal was then normalized to input signal density/area. Data shown is representative of three independent co-IP assays, averaged. The above graph is representative of three independent assays, averaged.

**Supplemental Figure 3. USP2a inhibition ablates proliferation in DG75 cells.** 1 million LMP1- or LMP1+ DG75 cells were treated with DMSO control, 10$\mu$M ML364, or 20$\mu$M ML364. At 24 hours, cells were counted with trypan blue to exclude dead cells. Statistics are comparing proliferation of each treatment group within the same cell line. Error bars represent standard deviation of three independent experiments. P values for significant differences (Student's t-test) are summarized by three asterisks ($p<0.001$), two asterisks ($p<0.01$), or one asterisk ($p<0.05$).
